# Comprehensive identification and characterization of conserved small ORFs in animals

**DOI:** 10.1101/017772

**Authors:** Sebastian D. Mackowiak, Henrik Zauber, Chris Bielow, Denise Thiel, Kamila Kutz, Lorenzo Calviello, Guido Mastrobuoni, Nikolaus Rajewsky, Stefan Kempa, Matthias Selbach, Benedikt Obermayer

## Abstract

There is increasing evidence that non-annotated short open reading frames (sORFs) can encode functional micropeptides, but computational identification remains challenging. We expand our published method and predict conserved sORFs in human, mouse, zebrafish, fruit fly and the nematode *C. elegans*. Isolating specific conservation signatures indicative of purifying selection on encoded amino acid sequence, we identify about 2000 novel sORFs in the untranslated regions of canonical mRNAs or on transcripts annotated as non-coding. Predicted sORFs show stronger conservation signatures than those identified in previous studies and are sometimes conserved over large evolutionary distances. Encoded peptides have little homology to known proteins and are enriched in disordered regions and short interaction motifs. Published ribosome profiling data indicate translation for more than 100 of novel sORFs, and mass spectrometry data gives peptidomic evidence for more than 70 novel candidates. We thus provide a catalog of conserved micropeptides for functional validation *in vivo*.

## Introduction

Ongoing efforts to comprehensively annotate the genomes of humans and other species revealed that a much larger fraction of the genome is transcribed than initially appreciated^1^. Pervasive transcription produces a number of novel classes of non-coding RNAs, in particular long intergenic non-coding RNAs (lincRNAs)^2^. The defining feature of lincRNAs is the lack of canonical open reading frames (ORFs), classified mainly by length, nucleotide sequence statistics, conservation signatures and similarity to known protein domains^2^. Although coding-independent RNA-level functions have been established for a growing number of lincRNAs^3,4^, there is little consensus about their general roles^5^. Moreover, the distinction between lincRNAs and mRNAs is not always clear-cut^6^, since many lincRNAs have short ORFs, which easily occur by chance in any stretch of nucleotide sequence. However, recent observations suggest that lincRNAs and other non-coding regions are often associated with ribosomes and sometimes in fact translated^7-16^. Indeed, some of the encoded peptides have been detected via mass spectrometry^10,17-23^. Small peptides have been marked as essential cellular components in bacteria^24^ and yeast^25^. More detailed functional studies have identified the well-known *tarsal-less* peptides in insects^26-29^, characterized a short secreted peptide as an important developmental signal in vertebrates^30^, and established a fundamental link between different animal micropeptides and cellular calcium uptake^31,32^.

Importantly, some ambiguity between coding and non-coding regions has been observed even on canonical mRNAs^15^: upstream ORFs (uORFs) in 5’ untranslated regions (5’UTRs) are frequent, well-known and mostly linked to the translational regulation of the main CDS^33,34^. To a lesser extent, mRNA 3’UTRs have also been found associated to ribosomes, which has been attributed to stop-codon read-through^35^, in other cases to delayed drop-off, translational regulation or ribosome recycling^36^, and even to the translation of 3’UTR ORFs (dORFs)^10^. Translational regulation could be the main role of these ORFs, and regulatory effects of translation (e.g., mRNA decay) could be a major function of lincRNA translation^12^. Alternatively, they could be ORFs in their own right, considering well-known examples of polycistronic transcripts in animals such as the *tarsal-less* mRNA^26-28^. Indeed, many non-annotated ORFs have been found to produce detectable peptides^10,17^, and might therefore encode functional micropeptides^37^.

Typically, lincRNAs are poorly conserved on the nucleotide level, and it is hard to computationally detect functional conservation despite sequence divergence even when it is suggested by synteny^2,38^. In contrast, many of the sORFs known to produce functional micropeptides display striking sequence conservation, highlighted by a characteristic depletion of nonsynonymous compared to synonymous mutations. This suggests purifying selection on the level of encoded peptide (rather than DNA or RNA) sequence. Also, the sequence conservation rarely extends far beyond the ORF itself, and an absence of insertions or deletions implies conservation of the reading frame. These features are well-known characteristics of canonical protein-coding genes and have in fact been used for many years in comparative genomics^39,40^. While many powerful computational methods to identify protein-coding regions are based on sequence statistics and suffer high false-positive rates for very short ORFs^41,42^, comparative genomics methods have gained statistical power over the last years given the vastly increased number of sequenced animal genomes.

Here, we present results of an integrated computational pipeline to identify conserved sORFs using comparative genomics. We greatly extended our previously published approach^10^ and applied it to the entire transcriptome of five animal species: human (*H. sapiens*), mouse (*M. musculus*), zebrafish (*D. rerio*), fruit fly (*D. melanogaster*), and the nematode *C. elegans*. Applying rigorous filtering criteria, we find a total of about 2000 novel conserved sORFs in lincRNAs as well as other regions of the transcriptome annotated as non-coding. By means of comparative and population genomics, we detect purifying selection on the encoded peptide sequence, suggesting that the detected sORFs, of which some are conserved over wide evolutionary distances, give rise to functional micropeptides. We compare our results to published catalogs of peptides from non-annotated regions, to sets of sORFs found to be translated using ribosome profiling, and to a number of computational sORF predictions. While there is often little overlap, we find in all cases consistently stronger conservation for our candidates, confirming the high stringency of our approach. Overall, predicted peptides have little homology to known proteins and are rich in disordered regions and peptide binding motifs which could mediate protein-protein interactions. Finally, we use published high-throughput datasets to analyze expression of their host transcripts, confirm translation of more than 100 novel sORFs using published ribosome profiling data, and mine in-house and published mass spectrometry datasets to support protein expression from more than 70 novel sORFs. Altogether, we provide a comprehensive catalog of conserved sORFs in animals to aid functional studies.

## Results

### Identification of conserved coding sORFs from multiple species alignments

Our approach, which is summarized in Fig. 1A, is a significant extension of our previously published method^10^. Like most other computational studies, we take an annotated transcriptome together with published lincRNA catalogs as a starting point. We chose the Ensembl annotation (v74), which is currently one of the most comprehensive ones, especially for the species considered here. In contrast to *de novo* genome-wide predictions^17,43^, we rely on annotated transcript structures including splice sites. We then identified canonical ORFs for each transcript, using the most upstream AUG for each stop codon; although use of non-canonical start codons has been frequently described^15-17,44,45^, there is currently no clear consensus how alternative translation start sites are selected. Next, ORFs were classified according to their location on lincRNAs or on transcripts from protein-coding loci: annotated ORFs serving as positive control; ORFs in 3’UTRs, 5’UTRs or overlapping with the annotated CDS; or on other transcripts from a protein-coding locus lacking the annotated CDS. We ignored pseudogene loci: although pseudogenes have been associated with a variety of biological functions^46-48^, their evolutionary history makes it unlikely that they harbor sORFs as independent functional units encoding micropeptides.

**Figure 1:**
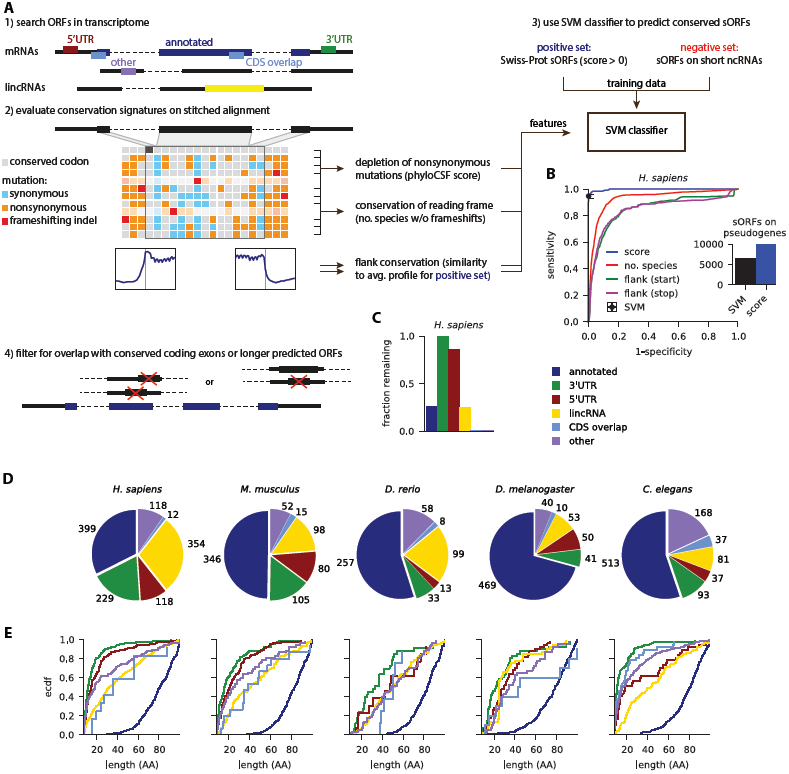
Identification of conserved sORFs in 5 animals. A) Overview of the pipeline. 1) Annotated transcripts are searched for ORFs and specific conservation features are extracted from the multiple species alignment (2). 3) A SVM classifier is used to predict coding sORFs (≤100 aa) with high specificity and sensitivity (B). 4) sORFs overlapping with larger predicted sORFs or with conserved annotated coding exons are removed (C). D) distribution of predicted sORFs in different regions of the transcriptome. E) length distribution of predicted sORFs.

Based on whole-genome multiple species alignments, we performed a conservation analysis to obtain four characteristic features for each ORF: most importantly, we scored the depletion of nonsynonymous mutations in the alignment using phyloCSF^49^; we also evaluated conservation of the reading frame from the number of species in the (un-stitched) alignment that lack frameshifting indels; finally, we analyzed the characteristic steps in nucleotide-level conservation (using phastCons) around the start and stop codons by comparing to the mean profile observed in annotated ORFs. Next, we trained a classifier based on support vector machines (see also Crappe et al.^50^ for a related approach) on confident sets of conserved small peptides and control sORFs from non-coding regions: as positive control, we chose conserved small peptides of at most 100 aa from Swiss-Prot with positive phyloCSF score. We discarded a number of presumably fast-evolving peptides: 177 in human and 77 in mouse, which are associated with antimicrobial defense, and 15 in fly of which 11 are signal peptides. As negative control, we chose sORFs on classical ncRNAs such as pre-miRNAs, rRNAs, tRNAs, snRNAs, or snoRNAs. Importantly, both of these sets overlap with a sizable number of genomic regions that are highly conserved on the nucleotide level (phastCons conserved elements; Fig. S1A). While each of the four conservation features performs well in discriminating positive and negative set (Fig. S1B), their combination in the SVM reaches very high sensitivity (between 1-5% false negative rate) and specificity (0.1-0.5% false positive rate) when cross-validating our training data (Fig. 1B and Fig. S1B). The classifier is dominated by the phyloCSF score (Fig. S1B), but the additional conservation features help to reject sORFs on annotated pseudogene transcripts, which typically do not show characteristic steps in nucleotide conservation near start or stop codons (Fig. 1B inset).

We noted that known small proteins typically reside in distinct genomic loci, while many predicted ORFs on different transcript isoforms overlap with one another or with annotated coding exons. Therefore, we aimed to remove candidates where the conservation signal could not be unambiguously assigned. We thus implemented a conservative overlap filter by excluding ORFs overlapping with conserved coding exons or with longer SVM-predicted ORFs (Methods). Most sORFs in 3’UTRs or 5’UTRs pass this filter, but many sORFs from different mRNA and lincRNA isoforms are collapsed, and most sORFs (85-99%) overlapping with annotated coding sequence are rejected (Fig. 1C and Fig. S1D).

### Hundreds of novel conserved sORFs, typically much smaller than known small proteins

With our stringent conservation and overlap filters, we predict 2002 novel conserved sORFs of 9 to 101 codons: 831 in *H. sapiens*, 350 in *M. musculus*, 211 in *D. rerio*, 194 in *D. melanogaster*, and 416 in *C. elegans*. Novel sORFs reside in lincRNAs and transcriptomic regions annotated as non-coding, with relatively few sORFs predicted in 3’UTRs or overlapping coding sequence relative to the size of these transcriptome regions (pre-overlap filter; see Fig. S1C). Our pipeline recovers known or recently discovered functional small peptides, such as all *tarsal-less* peptides^26-28^, sarcolamban^32^ and *pgc*^51^ in flies, *toddler*^30^ in zebrafish together with its human and mouse orthologs, and BRK1^52^ and myoregulin^31^ in human. We can confirm that many transcripts annotated as lincRNAs in fact code for proteins. However, it is a relatively small fraction (1-7%) that includes transcripts in intermediate categories, such as TUCPs in human^53^ and RITs in *C. elegans*^54^. Further, we note that a sizable number of uORFs are predicted to encode functional peptides, including the known case of MKKS^55^. Finally, we observe that the great majority of predicted sORFs is much smaller (median length 11 aa for 3’UTR sORFs in *C. elegans* to 49 aa for lincRNA sORFs in *D. rerio*) than annotated sORFs (median length 81-83 aa), with sORFs in 3’UTRs and 5’UTRs typically being among the shortest.

We assembled relevant information for the identified sORFs including coordinates, sequences, transcript models, and features analyzed in the following sections in Supplementary Tables 1-5.

### Novel sORFs are under purifying selection on the amino acid level

Since selection on the level of the encoded amino acid sequence permits synonymous sequence variation, we compared length-adjusted phyloCSF scores of predicted sORFs to those of control ORFs matched for their nucleotide-level conservation (Fig. 2A; methods). As expected from the design of our pipeline, we find that novel predicted sORFs are specifically depleted of nonsynonymous mutations, and in most cases to a similar extent as annotated ones. We also collected polymorphism data to perform a similar but independent test on a population genomics level: aggregating SNPs from all predicted sORFs (novel or annotated), we measured the dN/dS ratio and found that nonsynonymous SNPs are suppressed compared to synonymous ones to a greater extent than in control regions (Fig. 2B; Methods). This depletion is less pronounced than for annotated small proteins, and the associated *p*-values are lower in the species with higher SNP density (mouse and fruit fly with 16 and 45 SNPs/kb in the control regions) than in zebrafish or *C. elegans* with 1.7 and 2.0 SNPs/kb, respectively. It fails to pass the significance threshold in human with 2.4 SNPs/kb, where we get *p*=0.076 as the larger value from reciprocal X^2^tests.

**Figure 2:**
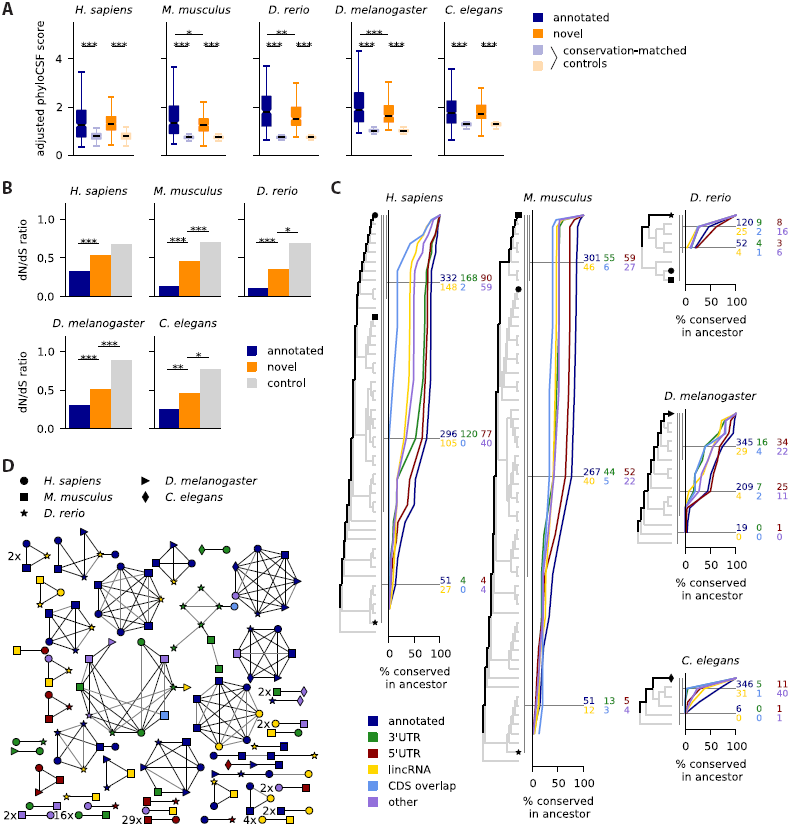
Predicted sORFs are under purifying selection and often widely conserved. A) Adjusted phyloCSF scores for predicted sORFs are higher than those from control sORFs matched by their nucleotide conservation level (phastCons). B) The dN/dS ratio of SNPs for novel predicted sORFs is smaller than for control ORFs in non-coding regions of the transcriptome, but larger than for annotated sORFs. C) Percentage of sORFs conserved in ancestral species as inferred from the multiple species alignment. Numbers for informative ancestors are indicated (e.g., the ancestors of primates, placental mammals and jawed vertebrates for *H. sapiens*). Symbols mark different reference species as in D). D) homology clustering of predicted sORFs in different species; only clusters with at least one non-annotated member and members from more than one species are shown, with multiplicity indicated. *** *p* < 0.001; ** *p* < 0.01; * *p* < 0.05; Mann-Whitney tests in A, reciprocal X^2^tests in B.

These results confirm that predicted sORFs permit synonymous more than nonsynonymous sequence variation when comparing within or between species, suggesting that selection acts on the level of the encoded peptide sequence and therefore implying functional peptide products.

### Some novel sORFs are widely conserved

We next sought to evaluate how widely the predicted sORFs are conserved. First, we took an alignment-based approach: we inferred most recent common ancestors from the alignment by tallying the species with conserved start and stop codons and (if applicable) splice sites, and without nonsense mutations. This analysis is dependent on the accuracy of the alignment, but it does not require transcript annotation in the aligned species. Using this method (see Fig. 2C) we find that after annotated small proteins, uORFs are most widely conserved, followed by the other sORF types. Of the novel sORFs found in human, 342 are conserved in placental mammals and 39 in the gnathostome ancestor (i.e., in jawed vertebrates). 18 are found conserved in teleosts, 49 in Drosophilids, and 88 in worms of the *Elegans* group.

We also addressed this question with a complementary analysis: we performed a homology clustering of sORFs predicted in the different species using a BLAST-based approach adapted for short amino acid sequences (Methods). This analysis clusters 1445 of in total 3986 sORFs into 413 homology groups, and 304 of 2002 novel predictions are grouped into 138 clusters. The clusters containing at least one novel predictions and sORFs from more than one species are summarily shown in Fig. 2D. Some novel predictions cluster together with sORFs annotated in other species, confirming the reliability of our approach and extending current transcriptome annotations. For instance, several zebrafish lincRNAs are found to encode known small proteins such as cortexin 2, nuclear protein transcriptional regular 1 (NUPR1), small VCP/p97-interacting protein (SVIP), or centromere protein W. Conversely, some lincRNAs from mouse and human encode small peptides with annotated (yet often uncharacterized) homologs in other species. Further, a sORF in the 3’UTR of murine Zkscan1 encodes a homolog of Sec61 gamma subunit in human, mouse, fish and fly. Also, a sORF in the 5’UTR of the worm gene *mnat-1* encodes a peptide with homology to murine *lyrm4* and the fly gene *bcn92*.

We also find 109 clusters of entirely novel predictions, such as 29 sORFs in 5’UTRs and 16 in 3’UTRs conserved between human and mouse, a 15 aa uORF in solute carrier family 6 member 8 (SLC6A8) conserved across vertebrates, or another 15 aa peptide from the 5’UTR of the human gene FAM13B conserved in the 5’UTRs of its vertebrate and fly homologs. One novel 25 aa peptide from annotated lincRNAs is predicted in three vertebrates and four other ones in two out of three. The other 22 human lincRNA sORFs found to be conserved in vertebrates (Fig. 2C) cluster together with annotated sORFs or are not detected in the other species for various reasons: they do not pass the overlap filter, do not use the most upstream start codon, or lack transcript annotation in mouse and zebrafish. Besides the 15 aa uORF peptide in FAM13B, there are also several peptides encoded in 3’UTRs or of mixed annotation conserved between vertebrates and invertebrates. Two clusters of unclear significance, consisting mainly of sORFs in the 3’UTRs of zinc-finger proteins, share a common HTGEK peptide motif, a known conserved linker sequence in C2H2 zinc fingers^56^. Finally, we note that our sequence-based approaches cannot resolve structural and/or functional homologies that persist despite substantial sequence divergence as observed between different animal peptides interacting with the Ca^2+^ ATPase SERCA^31,32^, or between bacterial homologs of the *E. coli* CydX protein^57^. We expect that further homologies between the predicted sORFs could be uncovered using more specialized approaches.

Taken together, this conservation analysis shows that novel sORFs are often widely conserved on the sequence level; further functional homologies could exist that are not detectable by sequence^31,32^.

### Conserved sORFs are predicted with high stringency

Many recent studies have addressed the challenge of identifying novel small protein-coding genes by means of computational methods or high-throughput experiments. These studies were performed in different species with different genome annotations, searching in different genomic regions, allowing different length ranges and using often quite different underlying hypotheses, for instance with respect to non-canonical start codons. Accordingly, they arrive at very different numbers. To reconcile these different approaches, we inclusively mapped sORFs defined in 15 other studies with published lists of coordinates, sequences or peptide fragments, to the comprehensive set of transcriptomic ORFs analyzed here (Supplementary Table 6). With the caveat that other studies often prioritized findings by different criteria, we then compared results with regard to the aspect of main interest here: conservation of the encoded peptide sequence, by means of comparative and population genomics as in Figs. 2A and B. We grouped studies by methodology, and by organism and genomic regions analyzed. We then compared sORFs predicted in our study but not in others to sORFs that were predicted elsewhere and analyzed but rejected here (Fig. 3). We used our results before applying the overlap filter. Considering changes in annotation (e.g., of coding sequences, lincRNAs and pseudogenes), we only compared to those sORFs that we analyzed and classified into the corresponding category. Generally, we find rather limited overlap between our predictions and results from other studies, which is partially explained by differences in applied technique and underlying hypothesis. We also find that the sORFs that we predict for the first time have consistently much higher length-adjusted phyloCSF scores than those found in other studies but rejected in ours; in many cases, we also find that the dN/dS ratio of nonsynonymous vs. synonymous SNP density is lower, albeit in a similar number of cases there is not enough data to render the *p*-value significant (we used the larger one from reciprocal X^2^-tests).

**Figure 3:**
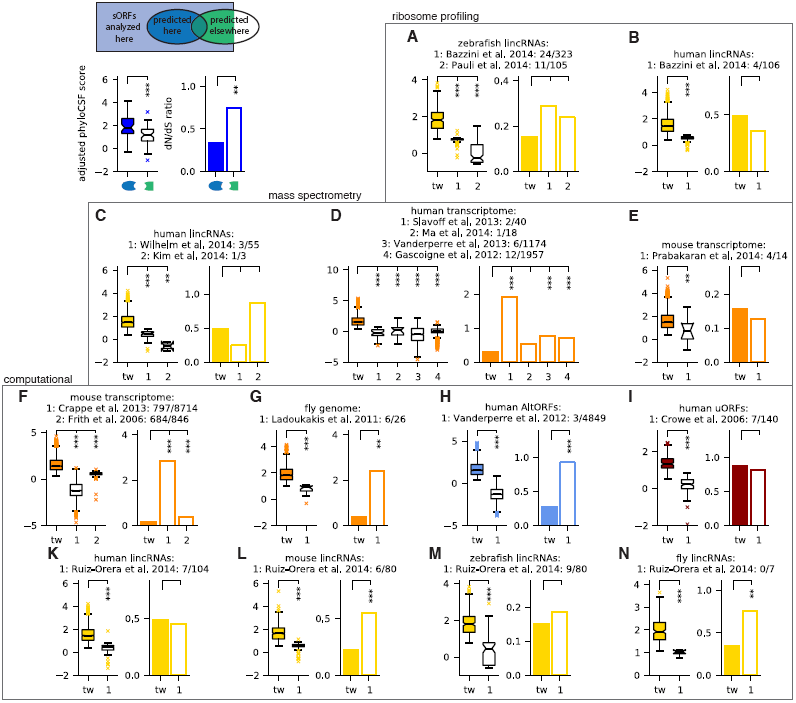
Predicted sORFs are under stronger selection than those found in other studies. Previous results obtained by ribosome profiling (A and B), mass spectrometry (C-E) or computationally (F-N) are compared with respect to their adjusted phyloCSF scores and the dN/dS ratio as indicated in the scheme (top left). For each publication analyzing sORFs in different organisms and genomic regions, the numbers of predicted sORFs that are also predicted here (before overlap filter) or at least analyzed, respectively, are given. phyloCSF scores and dN/dS ratios are compared for the sORFs that are predicted either here or in another study but not in both. tw: this work. *** *p* < 0.001; ** *p* < 0.01; * *p* < 0.05, using Mann-Whitney (phyloCSF scores) and reciprocal X^2^ tests (dN/dS), respectively.

First, we compared to a study using ribosome profiling in zebrafish^30^, with similar overlap as reported in our previous publication^10^, the results of which are re-analyzed with the updated transcriptome annotation for comparison (Fig. 3A-B). Ribosome profiling provides evidence of translation in the cell types or developmental stages analyzed, but in addition to coding sORFs it also detects sORFs with mainly regulatory functions such as uORFs. Next, we compared to 7 studies employing mass spectrometry^17-22^: matching given protein sequences or re-mapping detected peptides to the set of sORFs analyzed here, we find only between 1 and 12 common results from between 3 and almost 2000 sORFs (Fig. 3C-E). Note that up to 62% of peptides identified in these studies come from pseudogene loci which we excluded. While mass spectrometry provides direct evidence for peptide products, it is also performed in specific cell lines or tissues and has limited dynamic range. This can prevent detection of small peptides, which might be of low abundance or half-life, or get lost during sample preparation. Both experimental methods cannot distinguish sORFs coding for conserved micropeptides from those coding for lineage-specific or fast-evolving functional products. It is thus not surprising that these sORFs are as a group less conserved than the ones found using conservation as a selection criterion.

Next, we compared our results against other computational studies^18,43,50,58-60^. Here, we can often match much larger number of sORFs, but except for predictions of the CRITICA pipeline in mouse cDNAs^58^, we again find only limited overlap: we predict between 0 and 23% of analyzed sORFs found elsewhere, indicating a high variability in different computational methods, even though many of them use evolutionary conservation as a filter. The consistently better conservation indicators for our results (Fig. 3F-N) confirm that the deeper alignments and sensitive conservation features used here lead to increased performance. However, we remark that our method is not designed to find sORFs in alternative reading frames^61,62^ unless their evolutionary signal strongly exceeds what comes from the main CDS (e.g., because it is incorrectly annotated); also, the limited overlap with Ruiz-Orera et al.^60^ is not unexpected since their focus was on newly evolved lincRNA sORFs, which are by definition not well conserved. Finally, Crappé et al.^50^ and Ladoukakis et al.^43^ limited their search to single-exon sORFs, whereas 66% and 20% of sORFs predicted by us in the transcriptomes of mouse and fly, respectively, span more than one exon. However, even when restricting the comparison to single-exon sORFs, we find better conservation indicators for our results.

Given the consistently higher phyloCSF scores and often better dN/dS ratios of our sORFs when comparing to other studies, we conclude that our results present a high-stringency set of sORFs coding for putatively functional micropeptides.

### Novel peptides are often disordered and enriched for linear peptide motifs

We next investigated similarities and differences of sORF-encoded peptides to annotated proteins. First, we used amino acid and codon usage to cluster predicted sORFs, short and long annotated proteins and a negative control consisting of ORFs in non-coding transcriptome regions with small phyloCSF scores. Looking at amino acid usage, we were surprised to find that our novel predictions in four out of five species clustered with the negative control. However, when choosing subsamples of the data, novel predictions also often clustered together with annotated proteins, suggesting that their overall amino acid usage is intermediate. Indeed, the frequencies of most amino acids lie between those of positive and negative control. Interestingly, however, we found that novel predictions clustered robustly with annotated proteins when analyzing codon usage (with the exception of fruit fly).

Dissimilarity with annotated proteins was also confirmed when testing for homology to the known proteome using BLAST. Only a small fraction of novel predictions, mainly those in the ‘CDS overlap’ and ‘other’ categories, give significant hits (Fig. 4A). While some novel sORFs are homologous to annotated small proteins as revealed by the clustering analysis in Fig. 2C, there is no significant overlap between the sORFs that were assigned to homology clusters and those that have similarity to known proteins (Fisher’s *p* > 0.1 for all species except for *C. elegans* where *p*=0.003). Hence, even completely novel sORFs are sometimes conserved over wide distances.

**Figure 4:**
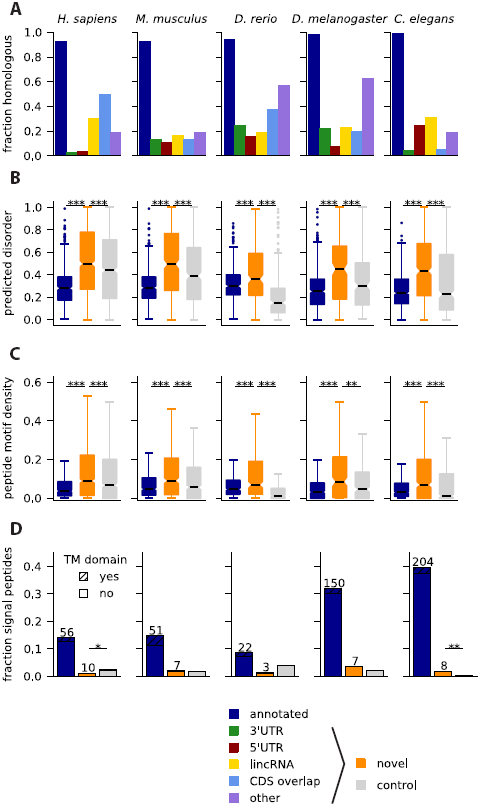
Properties of encoded peptide sequences. A) Only a small fraction of novel peptides has significant homology to known longer proteins. B) Novel predicted peptides are more disordered than annotated short proteins or conceptual products from length-matched control ORFs in non-coding regions, and they also have a higher density of linear peptide motifs (C). D) Some novel sORFs are predicted to encode signal peptides lacking trans-membrane (TM) domains, but not consistently more than expected. *** *p* < 0.001; ** *p* < 0.01; * *p* < 0.05, Mann-Whitney tests in B and C, binomial test in D.

We then hypothesized that differences in amino acid composition might give rise to different structural properties. We used IUPred^63^ to detect intrinsically unstructured regions, and found that novel predictions are much more disordered than known small proteins or a length-matched negative control (Fig. 4B). This could suggest that the peptides encoded by conserved sORFs adopt more stable structures only upon binding to other proteins, or else mediate protein-protein or protein-nucleic acid interactions^64^. It has recently become clear that linear peptide motifs, which are often found in disordered regions, can be important regulators of protein function and protein-protein interactions^65^. Indeed, when searching the disordered parts of sORF-encoded peptides for matches to motifs from the ELM database^66^, we find that the increased disorder comes with a higher density of such motifs in the predicted peptides (Fig. 4C), as was also observed recently for peptides identified with mass spectrometry^23^.

Since a recent study identified *toddler* and a number of other predicted signal peptides from non-annotated ORFs^30^, we searched our novel candidates with signalp^67^. Fig. 4D shows that a small number of our predicted sORFs have predicted signal sequences, and that most of these lack trans-membrane domains, but this does not exceed expectations from searching a length-matched control set. However, the typically lower amino acid conservation at the N-terminus of signal peptides could imply that some genuine candidates escape our conservation filters.

Taken together, these results show that novel sORF-encoded peptides are different from annotated proteins in terms of amino acid usage and sequence homology, that they are enriched in disordered regions and peptide motifs, and that only few of them encode signal peptides.

### 3’UTR sORFs are not consistently explained by stop-codon readthrough or alternative terminal exons

sORFs in 3’UTRs (dORFs) are least likely to be predicted as conserved compared to the other categories (Fig. S1C), but nevertheless we were surprised to find so many of them (between 33 in zebrafish and 229 in human). Although the existence of conserved dORFs was observed before^59^, and translation was also detected in ribosome profiling^10^, to the best of our knowledge there are no known examples of functional peptides produced from 3’UTRs (with the exception of known polycistronic transcripts). Therefore we explored the possibility that these ORFs actually represent conserved read-through events as suggested previously^35,68,69^, or come from non-annotated alternative C-terminal exons.

We first checked 283 read-through events in *Drosophila* previously predicted by conservation^68^, and 350 detected using ribosome profiling^35^. None of these coincides with any of the 41 sORF candidates we find in fly 3’UTRs, even though 3 of the candidates in Jungreis et al.^68^ were predicted as conserved and only rejected by the overlap filter. Similarly, none of 42 read-through events detected using ribosome profiling in human cells^35^ was predicted as conserved. However, three out of 8 known or predicted read-through events in human^70^ (in MPZ, OPRL1 and OPRK1) and one out of 5 read-through events predicted in *C. elegans* (in F38E11.6)^68^, were here incorrectly classified as 3’UTR sORFs (naturally, they have an in-frame methionine downstream of the annotated stop codon).

Given this small but finite number of false positives, we therefore explored our dORF candidates more systematically. In Fig. 4A, we had already established that dORF-encoded peptides have very little homology to known proteins, in contrast to the domain homology found in Drosophila readthrough regions^68^. Next, we checked that there is a very pronounced conservation step near the stop codon of annotated ORFs containing a predicted sORF in their 3’UTR, even though it is slightly smaller than for control ORFs lacking dORFs (Fig. 5A for human; see Fig. S5A for other species). This indicates that sequence downstream of the stop codons is indeed much less conserved and that these stops are not recently acquired (premature) stop codons or unused due to programmed frameshifts upstream. We made a number of further observations arguing against readthrough: dORFs are not generally close to the annotated stop codon or in the same frame, since we find only a small difference in the distribution of these distances and in most cases no preference for a specific reading frame (Fig. 5B and C; Fig. S5B and C); further, we observe a large number of intervening stop codons (Fig. 5D and Fig. S5D), and a step in conservation near the dORF start codons significantly more pronounced than for control ORFs in 3’UTRs (Fig. 5E and Fig. S5E). In addition, this observation makes it unlikely that dORFs represent non-annotated alternative terminal exons (where this methionine would not be associated with a conservation step).

**Figure 5:**
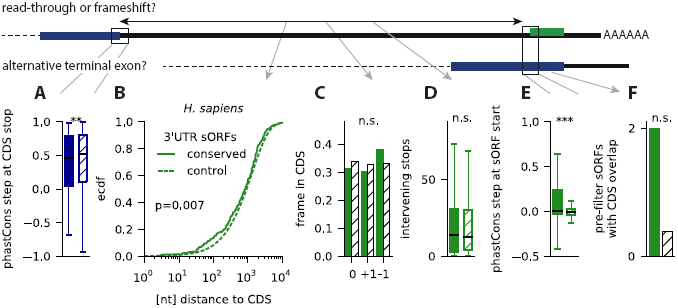
dORFs (sORFs in 3’UTRs) are not explained by stop-codon read-through or alternative terminal exons. Results are shown for *H. sapiens.* A) the step in the phastCons conservation track near the stop codon of the upstream CDS is only slightly less pronounced than for CDS without downstream conserved sORF. B) the dORFs are closer to the CDS than control sORFs, but they are not more often in the same frame (C), and they have a similarly high number of intervening in-frame stop codons (D). E) the step in the phastCons conservation track near start of predicted dORFs start is more pronounced than in other dORFs. F) Even before applying the overlap filter, very few predicted dORFs overlap with annotated coding exons. *** *p* < 0.001; ** *p* < 0.01; * *p* < 0.05; n.s. not significant. Mann-Whitney tests in A, D and E, Kolmogorov-Smirnov test in B, X^2^ test in C, Binomial test in F.

Further, if such un-annotated exons existed in large numbers, we would expect that at least some of our (pre-overlap filter) predictions overlap with already annotated alternative exons. However, except for *Drosophila* we only find at most two dORFs with CDS overlap, which is not more than expected compared to non-predicted dORFs (Fig. 5F and Fig. S5F).

In sum, these data suggest that our identification of 3’UTR sORFs is not systematically biased by conserved readthrough events or non-annotated terminal exons. Notably, we also identified candidates that clearly represent independent proteins, such as the dORF in the mouse gene Zkscan1 encoding a homolog of SEC61G, and a 22 aa dORF in the fly gene CG43200 which is likely another one of several ORFs in this polycistronic transcript.

### Experimental evidence for translation of and protein expression from predicted sORFs

Finally, we mined a large collection of publicly available and in-house generated data to verify translation and protein expression from predicted sORFs. In order to form expectations as to where and how highly our novel candidates could be expressed, we first analyzed publicly available RNA-seq expression datasets for different tissues (human and mouse) or developmental stages (zebrafish, fruit fly, and worm) (Supplementary Table 7). We then compared mRNAs coding for short proteins and lincRNAs with conserved sORFs with other mRNAs and lincRNAs, respectively (Fig. S6A). This analysis revealed that annotated short proteins come from transcripts with higher expression and lower tissue or stage specificity than long proteins. Conversely, it is well known that lincRNAs are not as highly and widely expressed as mRNAs^53,71^; we additionally find that lincRNAs with predicted sORFs are more highly and widely expressed than other lincRNAs. This analysis indicates that peptide products of novel sORFs could be of lower abundance than known small proteins, and that profiling translation or protein expression from a limited number of cell lines or tissues might not always yield sufficient evidence. We therefore used several datasets for the subsequent analysis.

First, we mined publicly available ribosome profiling datasets in various human and mouse tissues or cell lines, and in zebrafish (Supplementary Table 8). Several metrics to identify translated regions from such data have been proposed^9,14-16^; we rely here on the ORFscore method used in our previous publication^10^, which exploits the frame-specific bias of the 5’ positions of ribosome protected fragments to distinguish actively translated regions from those transiently associated with ribosomes or contaminants. It requires relatively deep coverage and a very clear 3 nt periodicity in ribosomal fragments, which is not always easily achievable (e.g., due to species-specific ribosome conformational properties^11,35^). We evaluated the ORFscore metric for datasets from human (HEK293 cells^45^, KOPT-K1 cells^72^ and human brain tissue^73^), mouse (embryonic stem cells^16^ and brain tissue^73^), and another zebrafish dataset^9^ in addition to the one used before^10^. The performance of these datasets was assessed by comparing ORFscore values of sORFs coding for annotated small proteins to those of the negative control from Fig. 1 by means of the Kolmogorov-Smirnoff *D* statistic; available datasets for *D. melanogaster*^35^ and *C. elegans*^74^ did not give a satisfying separation between positive and negative control (*D* < 0.55) and were not used.

Fig. 6A shows that predicted lincRNA sORFs have significantly higher ORFscores than the negative control (*p*-values between 2e-7 and 0.002), and similarly 5’UTR sORFs (*p*=2.5e-7 to 0.005) and sORFs in the “other” category (*p=*3.5e-7 to 0.04). sORFs in 3’UTRs reach marginal significance in some samples (*p*=0.02 for mouse brain and zebrafish). Choosing an ORFscore cutoff of 6 as done previously^10^, we find 45 novel sORFs translated in the human datasets, 15 in mouse, and 50 in zebrafish, respectively. We also find evidence for the translation of several non-conserved length-matched control sORFs, indicating that this set could contain lineage-specific or newly evolved coding ORFs or ORFs with regulatory functions.

**Figure 6:**
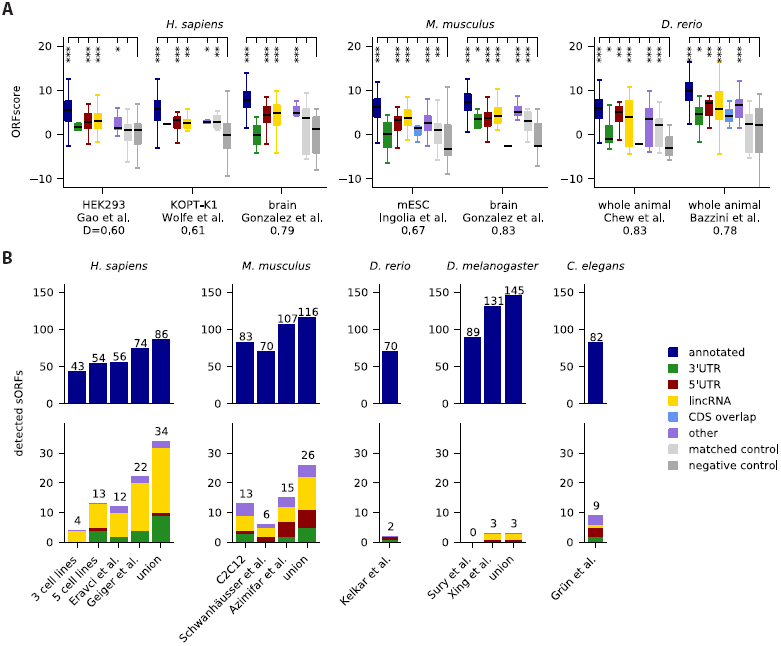
Experimental evidence supports translation of predicted sORFs and protein expression. A) Translation is detected using the ORFscore method10 on published ribosome profiling data. The Kolmogorov-Smirnov *D*-statistic is used to assess the performance of the dataset by comparing annotated sORFs to the negative control (dark gray). Length-matched sORFs from non-coding transcriptome regions are included as additional control (light gray). *** *p* < 0.001; ** *p* < 0.01; * *p* < 0.05 (Mann-Whitney test). B) Peptide expression of many predicted sORFs is confirmed by mining in house and published mass spectrometry datasets from cell lines and model organisms.

Next, we searched for peptide evidence in mass spectrometry datasets (Supplementary Table 9). We analyzed 3 in-house datasets to be published elsewhere: one for a mix of 3 human cell lines (HEK293, HeLa, and K562), one for a mix of 5 human cell lines (HepG2, MCF-10A, MDA-MB, MCF7 and WI38), and one for murine C2C12 myoblasts and myotubes. Further, we mined several published datasets: one for HEK293 cells^75^, one for 11 human cell lines^76^, one for mouse NIH3T3 cells^77^, one for mouse liver^78^, and whole-animal datasets from zebrafish^79^, fly^80,81^, and *C. elegans*^82^. All datasets were mapped with MaxQuant^83^ against a custom database containing our candidates together with protein sequences from UniProt. PSMs (peptide spectrum matches) were identified at 1% FDR, and those mapping to another sequence in UniProt with one mismatch or ambiguous amino acids were excluded. Using this strategy, we recover between 43 and 131 annotated small proteins per sample and confirm expression for 34 novel predictions in human, 26 in mouse, 2 in zebrafish, 3 in fly and 9 in *C. elegans* (Fig. 6B). For instance, we obtain PSMs for the recently described myoregulin micropeptide^31^ and for the long isoform of the fly *tarsal-less* gene^26-28^. In total, we find peptidomic evidence for 57 lincRNA sORFs. As observed previously in human^17,18^, mouse^23^ and zebrafish^10^ we also find PSMs for sORFs in 3’UTRs and 5’UTRs. MaxQuant output for PSMs and their supported sORFs is listed in Supplementary Tables 10-14, and the spectra with peak annotation are shown in Supplementary Figures 7-11.

In human and mouse, the results for novel predictions have considerable overlap of 17 and 8 hits, respectively, indicating that peptides from some sORFs can be reliably detected in multiple independent experiments. We also find more than one peptide for 9 and 11 novel sORFs in human and mouse and for one sORF in fly and worm, respectively. Likely as a consequence of the differences in expression on the RNA level (Fig. S6A), the PSMs supporting our novel predictions have generally lower intensities than those supporting the positive control (Mann-Whitney *p*=4e-9; Fig. S6C). However, we also observed that these PSMs are shorter than those mapping to UniProt proteins (*p=*0.005; Fig. S6D) and are of lower average quality: comparing Andromeda scores and other measures of PSM quality, we found that values for the PSMs supporting expression of novel predictions are smaller than for those mapping to the positive control (Fig. S6E-G). To test for the possibility of misidentifications, we therefore mapped two of our human datasets also against a 3-frame translation of the entire human transcriptome. As expected given the significantly (7.5fold) larger database, many PSMs (69 of 240) for annotated and novel sORFs now fall below the 1% FDR cutoff, but none of the spectra supporting the novel identifications is assigned to a different peptide sequence, and additional PSMs identified in these runs have similarly lower quality. Low-quality identifications can also result when posttranslational modifications of known proteins are not considered during the search^84,85^ (B. Bogdanov, H.Z. and M.S., under review). We therefore re-mapped one of the human datasets allowing for deamidation or methylation. Both possibilities again lead to a larger search space, such that 5 and 27 of 117 PSMs, respectively, fail to pass the FDR cutoff. Further, one of 14 PSMs supporting novel candidates is now attributed to a deamidated protein, but 7 of 103 PSMs mapping to sORFs in the positive control are also re-assigned, even though most of these sORFs have independent evidence from other PSMs. This suggests that targeted mass spectrometry approaches, complementary fragmentation techniques, or validation runs using synthetic peptides^23^ should be used to verify expression of ambiguous candidates. In summary, we cross-checked our predictions against a variety of high-throughput data: RNA-seq indicates that sORF-harboring lincRNAs are not as highly and widely expressed as other mRNAs, but more than lincRNAs without conserved sORFs. Analyzing ribosome profiling and mass-spectrometry data, we find evidence for translation and protein expression from 110 and 74 novel sORFs, respectively, across all datasets.

## Discussion

In our search for functional sORF-encoded peptides, we followed the idea that evolutionary conservation is a strong indicator for functionality if the conservation signal can be reliably separated from background noise and other confounding factors, such as overlapping coding sequences or pseudogenes. We therefore used conservation features that are very specific to known micropeptides (and canonical proteins), namely a depletion of nonsynonymous mutations, an absence of frameshifting indels, and characteristic steps in sequence conservation around start and stop codon. We then chose confident sets of positive and negative control sORFs, both of which have many members that are highly conserved on the nucleotide level, and combined these features into a machine learning framework with very high sensitivity and specificity.

Importantly, our refined pipeline also achieves a more reliable rejection of sORFs on pseudogene transcripts. Pseudogenes are important contaminants since frequent intervening stop codons imply that many of the resulting ORFs are short. While many pseudogenes are translated or under selective constraint,^48^ sORFs in these genes probably do not represent independent functional or evolutionary units.

Our integrated pipeline identifies sORFs comprehensively and with high accuracy, but we want to highlight a number of caveats and avenues for future research. First, the scope and quality of our predictions depends on the quality of the annotation: in some species, pseudogenes, lincRNAs and short ncRNAs (especially snoRNAs and snRNAs) have been characterized much more comprehensively, explaining some of the differences in the numbers seen in Fig. 1D. For instance, a recent study suggests that incomplete transcriptome assembly could lead to fragmented lincRNA identifications that obscure the presence of longer ORFs.^86^ Second, the quality of the predictions depends on the choice of the training data: while we aimed to choose negative controls that are transcribed into important RNA species and therefore often conserved on the nucleotide level, the training set is inevitably already separable by length alone, since there are only very few known small peptides below 50 aa, and very few ORFs on ncRNAs longer than that. A larger number of functionally validated very short ORFs would help to more confidently estimate prediction performance in this length range. Third, we remark that in some cases segmental duplications and/or genomic repeats give rise to a number of redundant sORFs, for instance in a 50kb region on zebrafish chromosome 9, or on chromosome U in flies. Fourth, our analysis is currently limited to finding canonical ORFs, even though usage of alternative initiation codons could be widespread^15-17,44,45^. Alternative start codon usage might even produce specific conservation signals that could be leveraged to confidently identify ORF boundaries.

Fifth, our approach is limited by the quality of the multiple species alignment: while the micropeptides characterized so far have very clear signatures allowing an alignment-based identification, there could be many instances where sequence conservation within the ORF and its flanking regions is not sufficient to provide robust anchors for a multiple alignment. For instance, functionally homologous micropeptides can be quite diverged on the sequence level. If additional homologous sequence regions can be reliably identified and aligned, a codon-aware re-alignment of candidate sequences^87^ could also help to improve detection power. Further, we currently only tested for a depletion of nonsynonymous mutations, but more sensitive tests could be implemented in a similar way^49^.

Sixth, since we did not find sORFs from our positive control or other known micropeptides to overlap with each other or longer ORFs, we used a quite conservative overlap filter to choose from each genomic locus one ORF most likely to represent an independent evolutionary and functional unit. This filter could be too restrictive: most importantly for sORFs overlapping annotated long ORFs in alternative reading frames, but also when the CDS annotation is incorrect, or for the hypothetical case that a micropeptide has multiple functional splice isoforms.

Finally, we specifically examined 3’UTR sORFs, for which mechanisms of translation are unclear. A very small number of cases could be explained by read-through or alternative exons, but we did not observe global biases. Depending on the experimental conditions, 3’UTR ribosome occupancy can be observed in *Drosophila* and human cells, but it has not been linked to active translation^36^. However, some mechanisms for downstream initiation have been proposed^88,89^, ribosome profiling gives evidence for dORF translation in zebrafish^10^, and some peptide products are found by mass-spectrometry^17-19,22,23^. Of course, the distinction between uORFs, main CDS, and dORFs becomes blurry for polycistronic transcripts.

To assess putative functionality of the encoded peptides, we tested our candidates for signatures of purifying selection; in addition to the expected depletion of nonsynonymous mutations in the multiple alignment when comparing to conservation-matched controls, we also found a weaker (but in many cases highly significant) depletion of nonsynonymous SNPs. A closer look at conservation statistics of identified sORFs revealed that many novel predictions are widely conserved between species (e.g., almost 350 in placental mammals and almost 40 in jawed vertebrates). By means of homology clustering, we observed that some of these novel predictions are actually homologous to known proteins, but we also found a sizable number of widely conserved uORFs and dORFs. Based on sequence homology, we could identify 6 novel predictions that are conserved between vertebrates and invertebrates. This small number is to be expected, since only two of 105 known annotated small proteins similarly conserved are shorter than 50 aa (OST4, a subunit of the oligosaccharyltransferase complex, and ribosomal protein L41), and only a minority of our predicted sORFs is longer than that (about 40% for zebrafish and 20% for the other species). Based on recently discovered functional and structural similarities between different SERCA-interacting micropeptides^31,32^, we expect that additional deep homologies between novel micropeptides might emerge in the future.

We also performed a systematic comparison to 15 previously published catalogs of sORF identifications, both computational and by means of high-throughput experiments. While underlying hypotheses, methods, and search criteria varied between studies, they shared the goal of extending genome annotations by identifying novel protein-coding regions. After matching results of other studies to our set of analyzed ORFs, we found in most cases quite limited overlap. However, we observed consistently better indicators of purifying selection for the set of sORFs identified here but not previously versus sORFs identified elsewhere but rejected here. This suggests that our conservative filters result in a high-confidence set of putatively functional sORFs, while a broad consensus about sORF characteristics has yet to emerge^37^. Most importantly, there could be a continuum between ORFs coding for micropeptides and those with regulatory functions (e.g., uORFs): we previously observed^90^ that several uORFs in *Drosophila* with regulatory functions controlled by dedicated re-initiation factors^89^ are also predicted here to encode putatively functional peptides, including the fly homolog of the uORF on the vertebrate gene FAM13B. A similar dual role could be fulfilled by sORFs on lincRNAs, whose translation could have the main or additional function of degrading the host transcript via nonsense-mediated decay^12^. Alternatively, such sORFs could represent evolutionary intermediates of novel proteins^60^.

Due to these and other ambiguities, a relatively limited overlap is not unexpected when combining computational and experimental approaches^10^: for instance, ribosome profiling provides a comprehensive snapshot of translated regions in the specific cell type, tissue and/or developmental stage analyzed. This includes sORFs that are translated for regulatory purposes or coding for fast-evolving or lineage-specific peptides such as the small proteins with negative phyloCSF scores excluded from our positive control set. A similar caveat applies to mass-spectrometry, which provides a more direct test of protein expression but has lower sensitivity than sequencing-based approaches, especially for low-molecular-weight peptides. The matching of measured spectra to peptide sequences is also nontrivial. Especially in deep datasets, low-quality PSMs can result from mismatched database hits if the database is incomplete or frequent post-translational modifications have not been considered^84,85^ (B. Bogdanov, H.Z. and M.S., under review).

Finally, we mined high-throughput RNA-seq, ribosome profiling and proteomics datasets to assess transcription, translation and protein expression of our predicted candidates. First, we used RNA-seq data to show that sORF-harboring lincRNAs are less highly and widely expressed than mRNAs (this is even more the case for lincRNAs without sORFs). In contrast, mRNAs with annotated sORFs are well and widely expressed, and in fact probably often encode house-keeping genes. Unfortunately, RNA expression is less useful as an expression proxy for the non-lincRNA categories due to an unknown translational coupling between main ORF and uORFs or dORFs. Given these findings, we expect that experiments for many different tissues, developmental time points, and environmental perturbations, and with very deep coverage, would be necessary to exhaustively profile sORF translation and expression. With currently available data, we could confirm translation of more than 100 conserved sORFs in several vertebrate ribosome profiling datasets using a stringent metric (ORFscore^10^), which exploits that actively translated regions lead to a pronounced 3 nt periodicity in the 5’ends of ribosome protected fragments. We also analyzed a number of published and in-house mass spectrometry datasets, and found peptidomic evidence for more than 70 novel candidates.

In conclusion, we present a comprehensive catalog of conserved sORFs in the transcriptomes of five animal species. In addition to recovering known small proteins, we discovered many sORFs in non-coding transcriptome regions. Most of these novel sORFs are very short and some are widely conserved between species. Based on the observation that encoded micropeptides are often disordered and rich in protein interaction motifs, we expect that they could function through protein-protein or protein-nucleic acid interactions. Given robust and confident signatures of purifying selection, and experimental evidence for translation and protein expression, our findings provide a confident starting point for functional analyses *in vivo*.

## Methods

### Transcriptome annotation and alignments

For all species, we used the transcript annotation from Ensembl (v74). Additionally, we used published lincRNA catalogs for human^53,91^, mouse^92^, zebrafish^38,93^ and fruit fly^94^, and added modENCODE^54^ transcripts for *C. elegans*.

We downloaded whole genome multiple species alignments from the UCSC genome browser (human: alignment of 45 vertebrates to hg19, Oct 2009; mouse: alignment of 59 vertebrates to mm10, Apr 2014; zebrafish: alignment of seven vertebrates to dr7, May 2011; fruit fly: alignment of 14 insect species to dm3, Dec 2006; worm: alignment of five nematodes to ce6, Jun 2008).

### ORF definition and classification

Spliced sequences for each transcript were scanned for the longest open reading frame starting with AUG and with a minimum length of at least 27 nucleotides. We scanned 4269 unstranded lincRNA transcripts from Young et al.^94^ on both strands. ORFs from different transcripts but with identical genomic coordinates and amino acid sequence were combined in groups and classified into different categories (using the first matching category for each group): “annotated” if an ORF was identical to the annotated coding sequence of a protein-coding transcript (i.e., biotype “protein coding”, and a coding sequence starting at the most upstream AUG, without selenocysteins, read-through or frameshift events). We classified ORFs as “pseudogene” if a member of a group came from a transcript or a gene locus annotated as pseudogene. We designated as “ncRNA” ORFs (negative controls) those with biotypes miRNA, rRNA, tRNA, snRNA or snoRNA. Next, “3’UTR” ORFs were classified as such if they resided within the 3’UTRs of canonical protein-coding transcripts, and if they did not overlap with annotated CDS (see below). Analogously, we assigned “5’UTR” ORFs. In the category “CDS overlap” we first collected ORFs that partially overlapped with 3’UTR or 5’UTR of canonical coding transcripts. ORFs in the “other” category were the remaining ones with gene biotype “protein coding”, or non-coding RNAs with biotypes “sense overlapping”, “nonsense-mediated decay”, “retained intron” or other types except “lincRNA”. Only those non-coding RNAs with gene and transcript biotype “lincRNA” were designated “lincRNA”. To exclude the possibility that alternative reading frames could be translated on transcripts lacking the annotated CDS, we finally added those ORFs that were completely contained in the annotated CDS of canonical transcripts to the “CDS overlap” category if other group members did not fall into the “other” category. Transcripts not from Ensembl were generally designated lincRNAs, except for *C. elegans*: in this case, we merged the modENCODE CDS annotation with Ensembl, and classified only the “RIT” transcripts as non-coding, while the ones that did not match the Ensembl CDS annotation were put in the “other” category. We then added Swiss-Prot and TrEMBL identifiers from the UniProt database (Nov 18 2014) to our ORFs by matching protein sequences.

### Predicting conserved sORFs using a SVM

From the multiple alignments for each ORF, we extracted the species with at least 50% sequence coverage and without frameshifting indels (using an insertion index prepared before stitching alignment blocks), recording their number as one feature. Stitched alignments for each putative sORF were then scored with PhyloCSF^49^ in the omega mode (options --strategy=omega -f6 –allScores) and the phylogenetic trees available at UCSC as additional input, yielding a second feature. Finally, we extracted phastCons conservation scores^95^ in 50 nt windows around start and stop codon (excluding introns but extending into flanking genomic sequence if necessary) and used the Euclidean distance of the phastCons profiles from the base-wise average over the positive set as third and fourth feature.

A linear support vector machine (LinearSVC implementation in the sklearn package in Python) was built using the four (whitened) conservation features and trained on positive and negative sets of sORFs. The positive set consisted of those sORFs in the “annotated” category with encoded peptide sequence listed in Swiss-Prot, with at most 100 aa (101 codons) length, some alignment coverage, and with positive phyloCSF score. The negative set consisted of sORFs from the “ncRNA” category with alignment coverage, but without overlap with annotated CDS.

We estimated the performance of the classifier by 100 re-sampling runs, where we chose training data from positive and negative set with 50% probability and predicted on the rest. Prediction of pseudogene sORFs (inset of Fig. 1B) was done either with the SVM, or based on the phyloCSF score alone, using a cutoff of 10 estimated from the minimum average error point in the ROC curve.

### Overlap filter

Refining our previous method, we designed an overlap filter as follows: in the first step, we only kept annotated sORFs or those that did not intersect with conserved coding exons. Here we took among the annotated coding exons in Ensembl (v74) or RefSeq (Sep 2 2014 for mouse, April 11 2014 for the other species) only those with conserved reading frame, requiring that the number of species without frameshifting indels reaches a threshold chosen from the minimum average error point in the ROC curves of Fig. 1B and S1 (11 species for human, 10 for mouse, 4 for zebrafish, 7 for fruit fly, and 2 for worm). In a second step we also required that the remaining ORFs were not contained in a longer ORF (choosing the longest one with the best phyloCSF score) that itself was predicted by the SVM and did not overlap with conserved coding exons. To exclude CDS overlap for the definition of 3’UTR and 5’UTR sORFs, and to design negative controls, we used Ensembl transcripts together with RefSeq (Feb 6 2014), and added FlyBase (Dec 12 2013) or modENCODE transcripts^54^ for fruit fly and worm, respectively (using intersectBed and a minimum overlap of 1bp between the ORF and CDS).

### Conservation analysis

For the analysis in Figs. 2A and 3, we computed adjusted phyloCSF scores as z-scores over the set of ORFs in the same percentile of the length distribution. Control ORFs were chosen among the non-annotated ORFs without CDS overlap and with their phyloCSF scores chosen among the 20% closest to zero and then sampled to obtain a statistically indistinguishable distribution of averaged phastCons profiles over the ORF.

SNPs were downloaded as gvf files from Ensembl (for human: v75, 1000 Genomes phase 1; for mouse, zebrafish and fly: v77); for *C. elegans* we took a list of polymorphisms between the Bristol and Hawaii strains from Vergara et al.^96^ and used liftOver to convert ce9 coordinates to ce6. We removed SNPs on the minus strand, SNPs falling into genomic repeats (using the RepeatMasker track from the UCSC genome browser, March 2015), and (if applicable) rare SNPs with derived allele frequency <1%. We then recorded for each ORF and its conceptual translation the number of synonymous and nonsynonymous SNPs, and the number of synonymous and nonsynonymous sites. For a set of sORFs, we aggregated these numbers and calculated the dN/dS ratio, where dN is the number of nonsynonymous SNPs per nonsynonymous site, and dS the number of synonymous SNPs per synonymous site, respectively. The control was chosen as before but without matching for nucleotide level conservation.

Alignment conservation in Fig. 2C was scored by analyzing for each ORF the multiple alignment with respect to the species where start and stop codons and (if applicable) splice sites were conserved, and where premature stop codons or frameshifting indels were absent. We then inferred the common ancestors of these species and plotted the fraction of ORFs with common ancestors at a certain distance to the reference species. For the homology graph in Fig. 2D we blasted sORF amino acid sequences from the different reference species against themselves and each other (blastp with options “-evalue 200000 -matrix PAM30 -word_size=2”). We then constructed a directed graph by including hits between sORFs of similar size (at most 20% deviation) for E-value < 10 and an effective percent identity PID_eff_ greater than a dynamically adjusted cutoff that required more sequence identity between shorter matches than longer ones (PID_eff_ = (percent identity) x (alignment length) / (query length); after inspecting paralogs or orthologs of known candidates such as *tarsal-less* and *toddler* we used the criterion PID_eff_ > 30+70 exp[-(query length + subject length)/20]). We then removed non-reciprocal edges, and constructed an undirected homology graph by first obtaining paralog clusters within species (connected components in the single-species subgraphs) and then adding edges for different reference species only for reciprocal best hits between paralog clusters. Finally, we removed singletons. For Fig. 2D, we combined isomorphic subgraphs (regarding sORFs in the same species and of the same type as equivalent), recorded their multiplicity, and plotted only the ones that contain sORFs from at least two different reference species and at least one novel prediction. For Fig. S1A we downloaded phastCons conserved elements from the UCSC genome browser (using vertebrate conserved elements for human, mouse and zebrafish; Nov 11 2014 for human and Nov 27 for the other species) and intersected with our set of ORFs; partial overlap means more than 50% but less than 99% on the nucleotide level.

### Comparison to other studies

We obtained results from other studies in different formats (Supplementary Table 6). Tryptic peptide sequences were mapped against the set of ORFs we analyzed (requiring preceding lysine or arginine). Amino acid sequences were directly matched to our set of ORFs, and ORF coordinates were matched to our coordinates (in some cases after conversion between genome versions or the removal of duplicate entries). Since different studies used different annotations and different length cutoffs, we then excluded from the matched ORFs the ones not in the category under consideration, e.g., longer ORFs, or sORFs that have since then been annotated or with host transcripts classified as pseudogenes. The remaining ones were compared to our set of predictions.

### Sequence analysis of encoded peptides

For Fig. 4A, we used blastp against the RefSeq database (Dec 2013) and collected among the hits with E-value > 10^-5^, percent identity > 50 and query coverage > 80% the best hit (based on percent identity) to entries of the same or larger length that were not flagged as “PREDICTED”, “hypothetical”, “unknown”, “uncharacterized”, or “putative”.

For the disorder prediction in Fig. 4B, we used IUPred^63^ in the “short” disorder mode and averaged disorder values over the sequence. For the motif discovery in Fig. 4C, we downloaded the file “elm_classes.tsv” from the ELM database website (http://elm.eu.org/downloads.html; Jan 27 2015). We then searched translated ORF sequences for sequences matches to any of the peptide motifs and kept those that fell into regions with average disorder > 0.5. For the signal peptide prediction in Fig. 4D, we used signalp v. 4.1^67^. Controls in Fig. 4B-D were chosen as in Fig. 2 but matched to the length distribution of novel predicted sORFs.

For Fig. S4A, we counted amino acid usage (excluding start and stop) for all ORFs; amino acids were sorted by their frequency in the positive control (“long ORFs”), which consists of annotated protein-coding ORFs from Swiss-Prot, whereas the negative control is the same as in Figs. 2 and 4 (not matched for conservation or length). We used hierarchical clustering with the correlation metric and average linkage on the frequency distribution for each group, and checked how often we obtained the same two clusters in 100 re-sampling runs where we took a random sample of ORFs in each group with 50% probability. For Fig. S4B, we counted codon usage, normalized by the amino acid usage, and then calculated a measure of codon bias for each amino acid using the Kullback-Leibler divergence between the observed distribution of codons per amino acid and a uniform one (in bits). We then performed clustering and bootstrapping as before.

### Analysis of 3’UTR sORFs

For all sORFs in the 3’UTR we obtained the annotated CDS of the respective transcript. We then computed the step in the phastCons conservation score (average over 25 nt inside minus average over 25 nt outside) at the stop codon of the annotated CDS and compared protein-coding transcripts with dORFs that are predicted and pass the overlap filter against other protein-coding transcripts. Similarly, we compared the step around the start codons of dORFs. We also compared the distance between the annotated stop and the start of the dORF, the distribution of the reading frame of the dORF start with respect to the annotated CDS, and the number of intervening stops in the frame of the annotated CDS. We finally checked how many predicted dORFs before applying the overlap filter overlap with annotated coding sequence and compared against the remaining dORFs.

We obtained read-through candidates from Supplementary Data 1 of Jungreis et al.^68^ and from Supplementary Tables 2 and 4 of Dunn et al.^35^ and matched the corresponding stop codons to stop codons in our set of 3’UTR sORFs.

### Expression analysis

For the expression analysis of sORF-containing transcripts we used RNA-seq data for 16 human tissues (Illumina Body Map), for 19 mouse tissues^97^, for 8 developmental stages of zebrafish^93^, 24 developmental stages for fruit fly from modENCODE and 8 developmental stages in *C. elegans*^82^ as shown in Supplementary Table 7. Reads were mapped to the reference genome using bowtie2 (with options --very-sensitive) except for human where we downloaded bam files from Ensembl; replicates were merged and then quantified using cufflinks and the Ensembl (v74) transcript annotation file together with the corresponding lincRNA catalogs. We ignored transcripts with all FPKM values below 10^-4^ and converted to TPM (transcripts per million^98^) as TPM= 10^6^ FPKM / (sum of all FPKM). Mean expression values were calculated by directly averaging TPM values of transcripts with nonzero TPM values over samples. Tissue or stage specificity was calculated as information content (IC) over the normalized distribution of transformed expression values 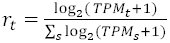 across tissues or stages, respectively, using the formula 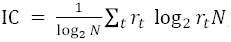 where *N* is the number of tissues or stages.

### Analysis of published ribosome profiling data

We obtained published ribosome profiling data as summarized in Supplementary Table 8. Sequencing reads were stripped from the adapter sequences with the Fastqx toolkit. The trimmed reads aligning to rRNA sequences were filtered out using bowtie. The remaining reads were aligned to the genome using STAR, allowing a maximum of 5 mismatches and ignoring reads that mapped to more than 10 different genomic locations. To reduce the effects of multi-mapping, alignments flagged as secondary alignments were filtered out. We then analyzed read phasing by aggregating 5’ read ends over 100 nt windows around start and stop of annotated coding sequences from Ensembl to assess dataset quality and obtain read lengths and 5’ offsets for use in scoring. From the datasets in Supplementary Table 8 we calculated the ORFscore as described previously^10^, pooling the reads from all samples if possible.

### Analysis of in-house and published mass spectrometry datasets

We used three in-house generated mass spectrometry datasets that will be published elsewhere: one in a mixture of HEK293, HeLa and K562 cells, one in a mixture of HepG2, MCF-10A, MDA-DB and MCF7 cells, and one in mouse C2C12 myoblasts and myotubes. Further, we mined published datasets in HEK293 cells from Eravci et al.^75^ 11 human cell lines from Geiger et al.^76^, in mouse NIH3T3 cells from Schwanhäusser et al. ^77^, in mouse liver from Azimifar et al. ^78^, in zebrafish from Kelkar et al.^79^, in flies from Sury et al.^81^ and Xing et al.^80^ and in *C. elegans* from Grün et al.^82^. All datasets (Supplementary Table 9) were searched individually with MaxQuant v1.4.1.2^83^ against a database containing the entire UniProt reference for that species (Swiss-Prot and TrEMBL; Nov 18 2014) merged with a database of common contaminant proteins and the set of predicted (annotated and novel) sORFs (after overlap filter). For fly datasets, an additional *E. coli* database was used. MaxQuant’s proteinFDR filter was disabled, while the peptide FDR remained at 1%. All other parameters were left at default values. To be conservative, we then remapped the identified peptide sequences against the combined database (treating Leucin and Isoleucin as identical and allowing for up to four ambiguous amino acids and one mismatch) with OpenMS^99^ and used only those peptides that uniquely mapped to our predictions. Features of PSMs (length, intensity, Andromeda score, intensity coverage and peak coverage) were extracted from MaxQuant’s msms.txt files. When re-mapping two human datasets (HEK293^75^ and 5 cell lines) against the 3-frame translation of the transcriptome, we created a custom database from all sequences longer than 7 aa between successive stop codons on transcripts from Ensembl v74 or published lincRNAs^53,91^. For the re-analysis of the HEK293 dataset^75^, we allowed deamidation (NQ) and methylation/methylester (KRCHKNQRIL) as additional variable modifications^85^.

## Acknowledgements

We thank the N Rajewsky lab for fruitful discussion, and Fabian Bindel for sharing unpublished mass-spec datasets. SDM and NR thank Francois Payre for initial discussions. SDM is funded by the Helmholtz-Alliance on Systems Biology (Max Delbrück Centrum Systems Biology Network), KK by the MDC-NYU exchange program, LC by the MDC PhD program, and BU through a Max-Delbrück fellowship of the MDC.

### Author contributions

NR and SDM initiated the project. SDM and BO designed and performed research for this paper. DT and BO performed conservation, sequence and expression analyses. SDM, LC and BO analyzed ribosome profiling data. HZ, CB, KK, GM and BO analyzed mass spectrometry data, supervised by SK and MS. BO prepared figures and wrote the paper, with input from the other authors..

### Competing interests

The authors declare no competing interests.

---

**Figure S1:**
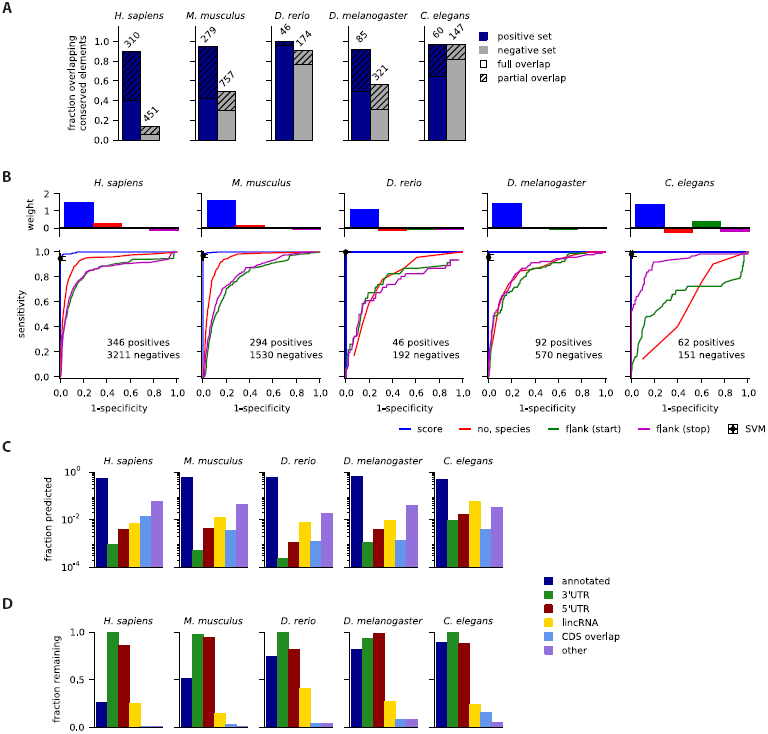
Overview of the pipeline (relating to Fig. 1). A) many sORFs from the positive control and from the negative control overlap fully or partially with phastCons conserved elements. B) The four conservation features all permit to separate positive from negative control (bottom panels); however, the phyloCSF score contributes most strongly to the SVM classifier. C) fraction of sORFs predicted as conserved (pre-overlap filter) for each category. D) fraction of sORFs retained after overlap filter in each category.

**Figure S4:**
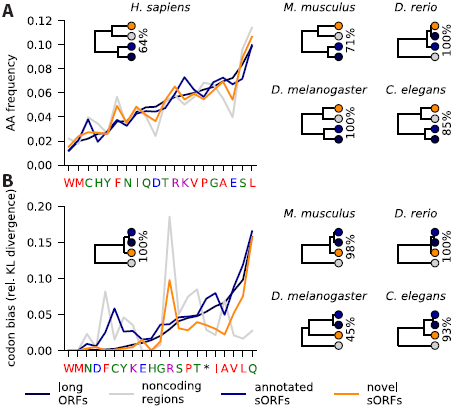
Sequence features of novel peptides (relating to Fig. 4). A) amino acid frequencies in long annotated ORFs, ORFs from noncoding control regions, predicted annotated sORFs and novel predicted sORFs are compared (shown for *H. sapiens*), and a hierarchical clustering is performed. Percentage values indicate how often the same clusters are obtained in a re-sampling analysis. Hydrophobic, acidic, basic and hydroxyl residues are colored red, blue, magenta and green, respectively. B) Codon bias is evaluated from the Kullback-Leibler divergence (Methods). Clustering done as in A.

**Figure S5:**
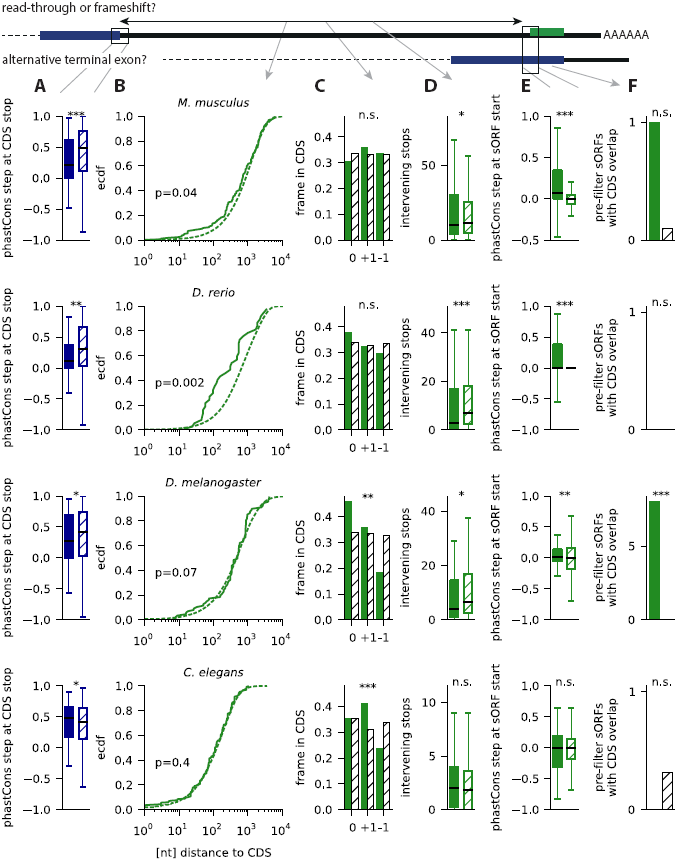
Properties of 3’UTR sORFs (same as Fig. 5 for the other species).

**Figure S6:**
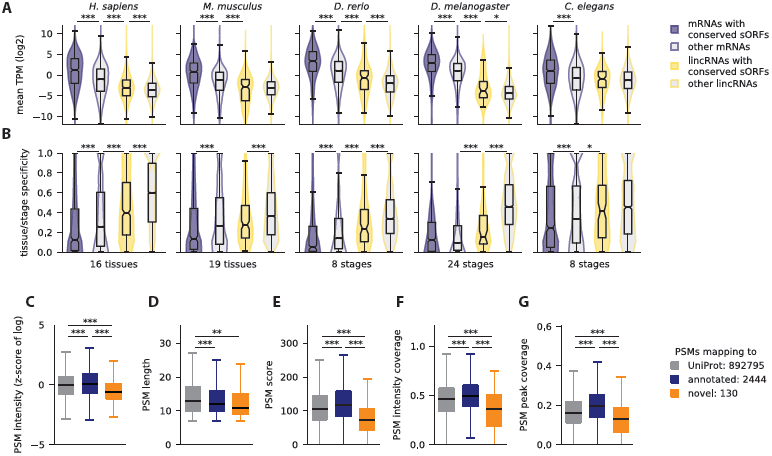
Expression analysis (relating to Fig. 6). A) violin and box plots of mean TPM values for mRNAs hosting predicted annotated sORFs, other mRNAs, lincRNAs hosting predicted novel sORFs and other lincRNAs, for 16 and 19 tissues in human and mouse, and 8, 24 and 8 developmental stages in zebrafish, fruit fly and *C. elegans,* respectively. B) violin and box plots of tissue or stage specificity for these transcripts. C) intensity for PSMs supporting annotated sORFs and peptides supporting novel predicted sORFs, aggregated over all datasets after log-transformation and normalization (z-score) relative to PSMs mapping to UniProt proteins. D) PSM length, E) Andromeda score, F) peak intensity coverage and G) peak coverage for PSMs as in C. *** p < 0.001; ** p < 0.01; * p < 0.05 (Mann-Whitney tests)

